# Common and lifestyle-specific traits of mycorrhiza-associated metabolite alterations in plant roots reflects strategies of root-mycorrhizal interactions

**DOI:** 10.1101/2022.01.27.478019

**Authors:** Mengxue Xia, Vidya Suseela, M. Luke McCormack, Peter G. Kennedy, Nishanth Tharayil

**Affiliations:** Department of Plant & Environmental Sciences, Clemson University, Clemson, SC 29634, USA; Center for Tree Science, The Morton Arboretum, Lisle, IL 60523, USA; Department of Plant and Microbial Biology, University of Minnesota, St Paul, MN 55108, USA

## Abstract

Convergent patterns in morphological and genetic traits of mycorrhizas have been well-documented and reflect common selection forces that define mycorrhizas. However, generalizable patterns of mycorrhiza-associated chemical alterations, which are immediately linked to plant and fungal strategies for successful symbiosis, have yet to be emerged. Comparing root metabolomes of phylogenetically-diverse plants inoculated by mycorrhizal fungi across two major lifestyles (arbuscular- *vs*. ecto-mycorrhizas), our study uncovers metabolite changes unique to each mycorrhizal lifestyle and those common across plant-mycorrhizal combinations irrespective of lifestyles. Arbuscular and ecto- mycorrhizal colonized roots accumulated different sets of carbohydrates, indicating unique carbon partitioning strategies: particularly, arbuscular mycorrhizal roots accumulated cyclic polyols inaccessible for symbionts, suggesting tighter regulation of plants in carbon partitioning. Mycorrhizas also altered specialized metabolism, featuring frequent increases of flavan-3-ols and decreases of flavanols irrespective of mycorrhizal lifestyles, suggesting tactical reconfiguration of specialized metabolites to facilitate/contain symbiosis. Our data show for the first time, to our knowledge, that part of the root metabolite alterations by mycorrhizas were relatively common across plant-mycorrhizal systems, highlighting their potentially critical regulatory and evolutionary role for successful symbiosis. This commonality appears robust to phylogenetic diversity of host plants and thus may be widespread in land plants. Our findings offer future research venues to elucidate the finer roles of these common traits of mycorrhiza-associated metabolite alterations and thus help to eventually develop a comprehensive understanding of this omnipresent plant-fungus partnership.

## Introduction

Most vascular plants (~92%) in modern terrestrial ecosystems form symbiosis with mycorrhizal fungi in their roots (Brundrett and Tedersoo, 2018). These widespread partnerships play a major role shaping terrestrial ecosystems as they mediate plant nutrition, plant responses to the environment, and ecosystem biogeochemical cycles (Brundrett and Tedersoo, 2018; Genre et al., 2020). Thus, knowledge on the biological processes underlying plant-mycorrhizal interactions is essential to the stewardship of ecosystem services provided by mycorrhizas, such as plant productivity, crop quality, and ecosystem carbon balance. One common theme of plant-mycorrhizal interactions is convergent evolution (Strullu-Derrien et al., 2018; Genre et al., 2020; Miyauchi et al., 2020). About 50,000 soil fungal taxa with highly diverse evolutionary history have converged to four main mycorrhizal lifestyles, among which arbuscular (AM) and ecto-mycorrhizal (EM) fungi dominate most terrestrial ecosystems (Brundrett and Tedersoo, 2018; Genre et al., 2020). EM independently evolved up to 80 times from multiple lineages spanning hundreds of millions of years but have converged to exhibit dimorphic root structures, intercellular Hartig network, and a thin yet strengthened cortex, in contrast to AM fungi that show less effect on root morphology and form intracellular arbuscles (Brundrett, 2002; Tedersoo and Smith, 2017). On the genetic level, AM and EM shared common traits such as lack/loss of lignin-cellulose degrading enzymes and expansion of functionally similar small-secreted protein families, some of which have been shown to counteract plant defense (Kloppholz et al., 2011; Plett et al., 2014). These convergent patterns within or across mycorrhizal lifestyles in morphological and genetic traits of mycorrhizas have been recently reviewed (Genre et al., 2020) and reflect common selection forces that define mycorrhizal symbiosis.

However, the common *vs*. lifestyle-specific traits of mycorrhiza-associated alterations on plant metabolomes that directly facilitate/ensue mycorrhizas remain largely unexplored. Compared to genomic and transcriptomic traits, plant metabolomes represent the intermediates/end products of cellular regulation, are closer to biological functions, and reflect immediate responses of plants to their environments (Fiehn et al., 2002; Feussner and Polle, 2015; Alseekh et al., 2020); thus, plant metabolomes would also face direct selection pressures to sustain a beneficial partnership with mycorrhizal fungi and may be subject to convergent alterations in response to the similar challenge for maintaining mycorrhizas. Plant-mycorrhizal interactions have been shown to reprogram plant metabolomes as a prerequisite for the establishment of functional mycorrhizas (*e.g*., Ouledali et al., 2018, carbohydrates; Salvioli et al., 2012, nitrogen-containing metabolites; Szuba et al., 2020, specialized metabolites), by shaping the transport/partitioning of photosynthates and mineral resources and modulating plant defense (Schweiger et al. 2014; Genre et al., 2020). Nevertheless, comparison of mycorrhiza-associated metabolite alterations across plant/fungal species or mycorrhizal lifestyles proves elusive, partly because plant metabolites are highly diversified while metabolite studies often focused on different subsets of metabolites of interest. A seminal study compared metabolite alterations in multiple plant species that faced the identical challenge of sustaining mycorrhizas with AM fungus *Rhizophagus irregularis* (Schweiger et al., 2014). Surprisingly, this study found little commonality but high-level plant-species specificity, with important implication that transferability of metabolite alterations observed from a few model species to other plants is limited. However, this work investigated leaf metabolites. Leaves have been shown to be less affected by mycorrhizas than roots that directly accommodate fungal partners (Xia et al., 2021). Taken together, although being a crucial piece of the puzzle for the biological processes underlying plant-mycorrhizal interactions/strategies to maintain mycorrhizas, commonality *vs*. specificity of metabolite alterations in plant roots remained unexplored. We initiate this study to compare mycorrhiza-associated metabolite alterations in roots across various combination of plant hosts and mycorrhizal fungal species, which would facilitate the transferability of knowledge from model species to broader plant-mycorrhizal systems.

We first ask if distinct metabolite alterations would align with unique mycorrhizal lifestyles. AM and EM have been shown to be associated with contrasting plant functions and ecosystem biogeochemical processes (*e.g*., Phillips et al., 2013; Soudzilovskaia et al., 2015; Chen et al., 2016). Understanding how these different lifestyles may differentially modulate plant metabolome would reveal finer-level mechanisms underlying their respective roles on plant and ecosystem functioning. Prior studies suggest that AM *vs*. EM may have divergent impacts on root carbohydrate profiles, as they tend to exhibit differential carbon (C) partitioning patterns. EM symbionts often require more C than AM symbionts (van der Heijden et al., 2015). In EM-dominated plant communities within a subarctic-alpine ecosystem, the estimated C pool in EM mycelium was similar to that stored in plant biomass; by contrast, AM mycelium retained much less C (<25%) compared to plant biomass C in AM-dominated communities (Soudzilovskaia *et al*., 2015). Isotope evidence that estimated allocation of photosynthates was variable but suggest that < 10% photosynthetically fixed C was typically allocated to AM mycelium (Jakobsen and Rosendahl, 1990; Johnson et al., 2002ab) while EM could acquire 10 to 50% fixed C (Rosling et al., 2004; Hobbie, 2006) and drive C flux between host plants (Simard et al, 1997). AM and EM also showed differential reciprocal relationships between the C invested by plants and the nutrients returned by fungi in a multiple-species comparative study (Keller and Phillips, 2019): in AM-associated plants, greater C investment belowground was linked with higher nitrogen (N) uptake, whereas in EM plants, this positive relationship was not observed, and high C investment often corresponded to low N return. These observations suggest that EM and AM may differ in their C partitioning patterns: EM symbiosis could be a stronger C sink, while the transfer of C from plant hosts to AM fungi may be more under plant control. This functional difference may reflect on the carbohydrate profiles in mycorrhizal roots, as fine-tuning of carbohydrates often represents a means to mediate C partitioning between plants and their fungal partners. For example, EM fungi in functional symbiosis quickly transformed hexoses to large quantities of fungal carbohydrates such as mannitols (up to >60% of total fungal carbohydrates, Lewis and Harley, 1965; Ceccaroli et al., 2003; Nehls et al., 2007; Plett et al., 2020). This removal of hexoses may help maintain a sharp gradient at the exchange site and thus facilitate C efflux (mainly through a concentration-dependent transporter family, An et al., 2019) from plants to mycorrhizal fungi. Accordingly, we hypothesize that (**1**) AM and EM associations would differentially alter root carbohydrate profiles, reflecting their distinct C partitioning strategies. We predicted that, compared to their respective non-inoculated controls, the carbohydrate profiles in EM-colonized roots may maintain a greater proportional abundance of fungal-specific carbohydrates than AM roots, thereby maintaining a continuous, strong C sink.

We next ask if there are common metabolite alterations that frequently occur in plant-mycorrhizal systems irrespective of mycorrhizal lifestyles, reflecting similar selection forces shaping plant-mycorrhizal interactions. A common evolutionary trend of mycorrhizas across AM and EM is the expression of the functionally similar genes (*e.g*., nutrient transporters and small secreted proteins), where the small secreted proteins could dampen plant defense responses (Genre et al., 2020). Consistently, both AM and EM have been shown to elicit only transient transcriptomic responses linked to plant defense/stress responses, which were later attenuated in established symbiosis (Liu et al., 2003; Tarkka et al., 2013; Giovannetti et al., 2015). If these genetic/transcriptional traits could be translated to chemistry/metabolite alterations, one would expect no accumulation or lower-level of defense/stress related specialized compounds in established mycorrhizal roots. This is supported by observations that the establishment of mycorrhizas were associated with decreased or unaffected levels of protective compounds in roots (*e.g*., cell-wall bound ferulic acids, tannins, Münzenberger et al., 1995; Solaiman et al., 2018; Sebastiana et al., 2021) and lack of additional lignin deposition (Chen et al., 2021). On the other hand, mycorrhizas have been shown to induce substantial accumulation of phenolic compounds in the endodermis or exodermis of root cortex (Ling-Lee et al., 1977; Weiss et al., 1997, 1999), with these specific root compartments proposed crucial in the evolution of mycorrhizas by forming a barrier to detrimental fungal penetration (Brundrett, 2002; Strullu-Derrien et al., 2009). These seemingly contradictory patterns reflect a common requisite for mycorrhization (*sensu* Brundrett, 2002): accommodating mycorrhizas while protecting root tissues from detrimental fungal growth (*e.g*., penetration to vascular tissues). This general selection along plant-mycorrhizal interactions may result in convergent metabolite modulation. Accordingly, we hypothesize that (**2**) the mycorrhiza-associated metabolite alternations in roots would exhibit common patterns across AM and EM lifestyles, reflecting chemical events critical for root-mycorrhizal interactions. Specifically, the suppression of a subset of root defense compounds to facilitate symbiosis would be accompanied by an upregulation of a different subset of specialized compounds which could be important for protecting root tissues in the presence of fungal colonization.

Here, we test these hypotheses by examining mycorrhiza-induced modulation of root metabolites in multiple plant-mycorrhizal combinations across AM and EM lifestyles in a phylogenetically-diverse set of temperate tree species. Using multi-species comparison and comprehensive metabolomics approaches, our data uncovered novel chemical events unique to or shared across AM and EM lifestyles, with implications for carbon partitioning strategies and tissue protection that could be critical for maintaining mycorrhizas.

## Results

### Mycorrhizal colonization, plant growth, and nutrient status

Both AM fungi *Gigaspora margarita* (Gm) and *Funneliformis mosseae* (Fm) colonized their hosts *Juniperus virginiana* (JUVI) and *Liriodendron tulipifera* (LITU) at high and consistent levels (67.6 – 75.4%, Table 1). Colonization by EM was more variable, typically ranging from 30 - 75% depending on the fungal identity. Mycorrhizal colonization improved plant growth and/or the levels of N (gauged by the ratio of carbon and nitrogen, C/N) in plant tissues across all plant-fungus combinations, indicating functional symbiosis. Specifically, the seedlings of JUVI, LITU, and *Pinus taeda* (PITA) exhibited improved growth by two mycorrhizal fungal species (Table 1). Although colonization by Ss and *Tuber sp*. (Ts) did not result in a better growth for PITA and *Quercus macrocarpa* (QUMA), respectively, it led to a lower C/N (thus higher tissue N levels) in either their leaves or roots (Table 1). Because metabolites may also depend on plant growth and nutrient utilization, we performed variance partitioning analysis to compare the potential drivers (*i.e*., mycorrhizal colonization, plant growth, and/or nitrogen status) of metabolite responses in the following analysis.

**Table 1.**
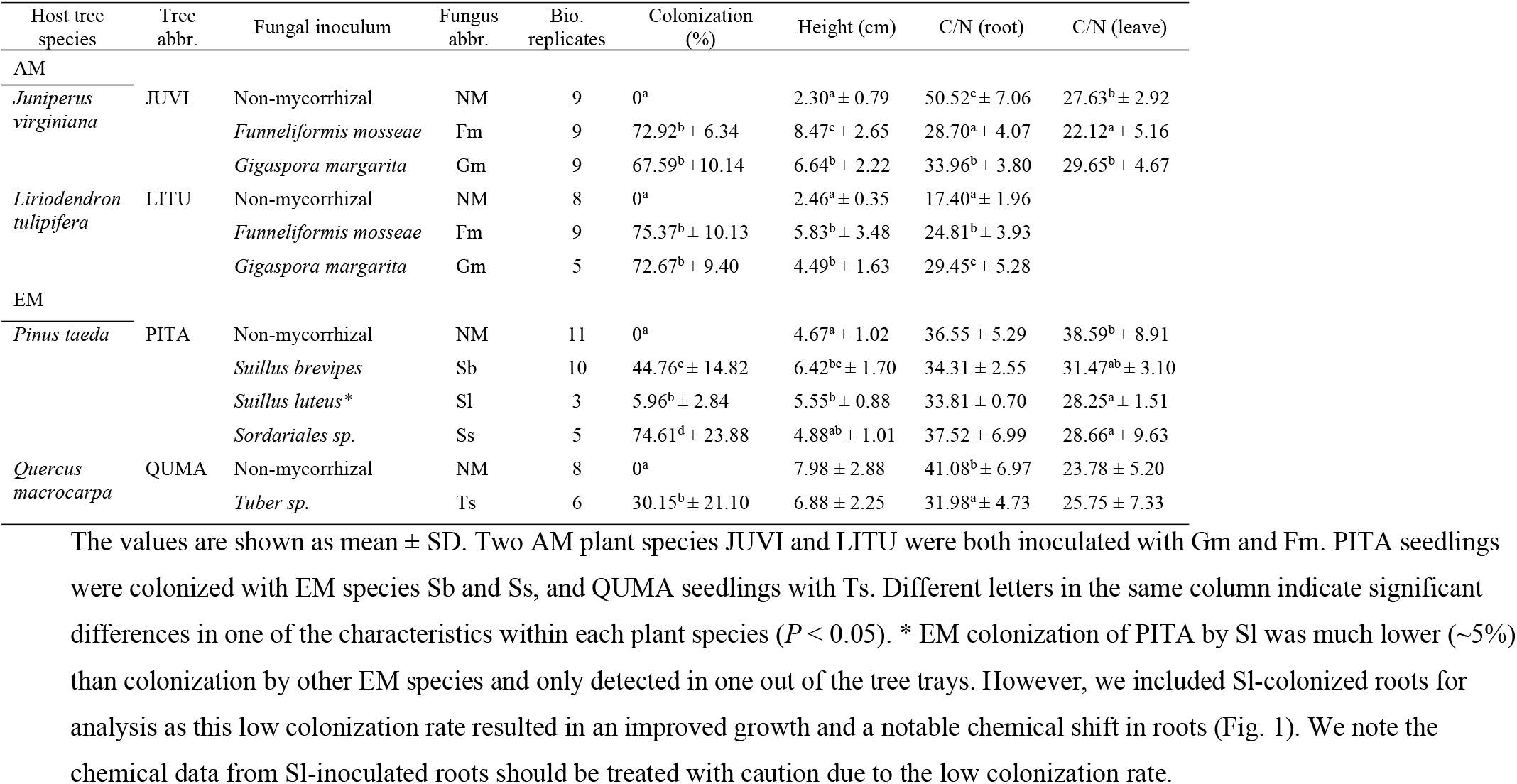
Mycorrhizal colonization rate (%), seedling height, and the ratios of carbon and nitrogen (C/N) of seedings inoculated with AM or EM fungi.

| Host tree species | Tree abbr. | Fungal inoculum | Fungus abbr. | Bio. replicates | Colonization (%) | Height (cm) | C/N (root) | C/N (leave) |
| --- | --- | --- | --- | --- | --- | --- | --- | --- |
| AM |  |  |  |  |  |  |  |  |
| *Juniperus virginiana* | JUVI | Non-mycorrhizal | NM | 9 | 0<sup>a</sup> | 2.30<sup>a</sup> ± 0.79 | 50.52<sup>c</sup> ± 7.06 | 27.63<sup>b</sup> ± 2.92 |
|  |  | *Funneliformis mosseae* | Fm | 9 | 72.92<sup>b</sup> ± 6.34 | 8.47<sup>c</sup> ± 2.65 | 28.70<sup>a</sup> ± 4.07 | 22.12<sup>a</sup> ± 5.16 |
|  |  | *Gigaspora margarita* | Gm | 9 | 67.59<sup>b</sup> ± 10.14 | 6.64<sup>b</sup> ± 2.22 | 33.96<sup>b</sup> ± 3.80 | 29.65<sup>b</sup> ± 4.67 |
| *Liriodendron tulipifera* | LITU | Non-mycorrhizal | NM | 8 | 0<sup>a</sup> | 2.46<sup>a</sup> ± 0.35 | 17.40<sup>a</sup> ± 1.96 |  |
|  |  | *Funneliformis mosseae* | Fm | 9 | 75.37<sup>b</sup> ± 10.13 | 5.83<sup>b</sup> ± 3.48 | 24.81<sup>b</sup> ± 3.93 |  |
|  |  | *Gigaspora margarita* | Gm | 5 | 72.67<sup>b</sup> ± 9.40 | 4.49<sup>b</sup> ± 1.63 | 29.45<sup>c</sup> ± 5.28 |  |
| EM |  |  |  |  |  |  |  |  |
| *Pinus taeda* | PITA | Non-mycorrhizal | NM | 11 | 0<sup>a</sup> | 4.67<sup>a</sup> ± 1.02 | 36.55 ± 5.29 | 38.59<sup>b</sup> ± 8.91 |
|  |  | *Suillus brevipes* | Sb | 10 | 44.76<sup>c</sup> ± 14.82 | 6.42<sup>bc</sup> ± 1.70 | 34.31 ± 2.55 | 31.47<sup>ab</sup> ± 3.10 |
|  |  | *Suillus luteus* * | Sl | 3 | 5.96<sup>b</sup> ± 2.84 | 5.55<sup>b</sup> ± 0.88 | 33.81 ± 0.70 | 28.25<sup>a</sup> ± 1.51 |
|  |  | *Sordariales sp.* | Ss | 5 | 74.61<sup>d</sup> ± 23.88 | 4.88<sup>ab</sup> ± 1.01 | 37.52 ± 6.99 | 28.66<sup>a</sup> ± 9.63 |
| *Quercus macrocarpa* | QUMA | Non-mycorrhizal | NM | 8 | 0<sup>a</sup> | 7.98 ± 2.88 | 41.08<sup>b</sup> ± 6.97 | 23.78 ± 5.20 |
|  |  | *Tuber sp.* | Ts | 6 | 30.15<sup>b</sup> ± 21.10 | 6.88 ± 2.25 | 31.98<sup>a</sup> ± 4.73 | 25.75 ± 7.33 |

### Global changes of root metabolome induced by mycorrhizas

Mycorrhizal colonization reconfigured root metabolomes in all plant-fungus combinations across AM and EM (Fig. 1). A large number of individual mass features were extracted from root tissues, ranging from 6920 features for PITA roots to 10405 features for JUVI roots, where a substantial part of these features were altered by mycorrhization. The differentially regulated features by mycorrhizas (Student’s t-test, *P* < 0.05) were visualized in *van* Krevelen diagrams (Fig. 1A). AM differentially regulated 4494 (LITU×Gm) to 7231 (JUVI×Fm) features that span all major chemical classes, while EM altered 1288 to 3439 features (Fig. 1A). This large difference in the number of altered features between AM and EM were less likely due to colonization rate. For example, AM fungus Fm and EM fungus Ss exhibited similar colonization rates (~70%) in LITU and PITA, respectively (Table 1); however, Fm significantly altered four-fold more features in LITU than Ss in PT (5270 *vs*. 1288, Fig. 1A).

**Fig. 1.**
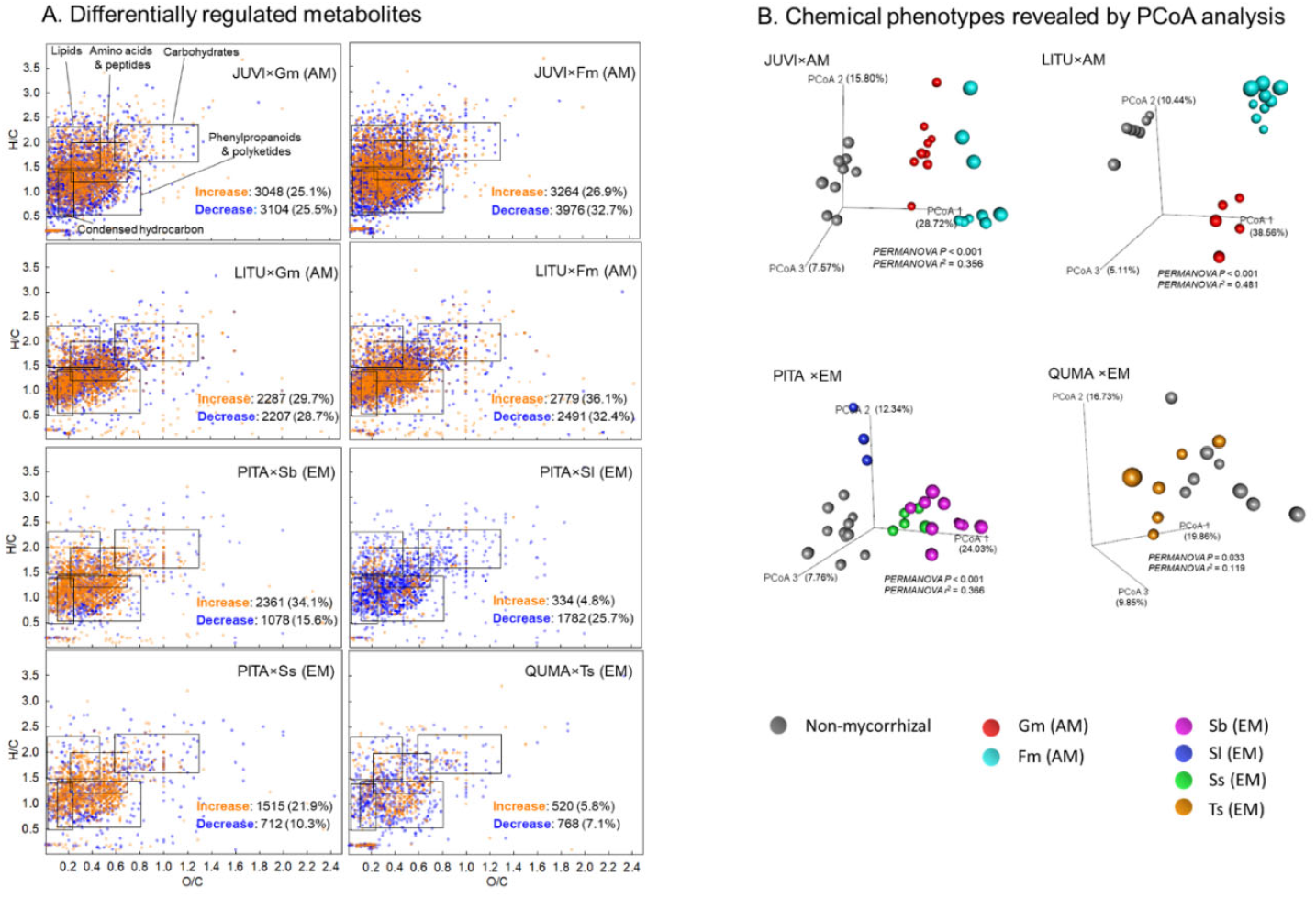
Comparing chemical landscapes between non-mycorrhizal and mycorrhizal roots in eight plant-mycorrhizal fungus combinations across AM and EM lifestyles. **A.** *van* Krevelen diagrams show the distribution of differentially regulated metabolites (two-tailed Student’s *t*-test, *P* < 0.05) in major chemical classes. The total numbers of increased and decreased metabolites by mycorrhization were shown. **B.** Principal coordinates analysis (PCoA) shows the separation of root chemical phenotypes between individual non-mycorrhizal seedlings and seedlings colonized by different mycorrhizal fungal species. The significance of separation is indicated by PERMANOVA *P* values, while the effect size is indicated by PERMANOVA *r^2^* values.

Taking into account the entire chemical landscapes, principal coordinates analysis (PCoA) showed that the four plant species separated into four distinct chemical phenotypes, with two gymnosperms closer to each other and further away from the two angiosperms, mirroring their phylogenetic relationships (Fig. S1). Within each plant species, colonized roots exhibited significantly different chemical phenotypes from those uncolonized (PERMANOVA *P* < 0.033, PERMANOVA *r*^2^ ranges from 0.119 to 0.481, Fig. 1B). Not only did the chemical phenotypes differ between colonized *vs*. uncolonized roots, roots from the same species but colonized by different fungal species also separated into distinct clusters (Fig. 1B), indicating that the effect of mycorrhizas on root metabolomes were significant and specific to fungal species. Together, these patterns of unannotated mass features demonstrated significant global metabolic alterations by mycorrhizas in all plant-fungus combinations included in this study and set the stage for understanding common *vs*. unique metabolite responses across various plant-mycorrhizal systems.

### Alterations of primary metabolism

Mycorrhizas affected most major groups of primary metabolites including carbohydrates (further grouped as disaccharides, pentoses, hexoses, and polyols, a carbohydrate derivative), amino acids, and organic acids (Fig. 2A, S2). AM associations and the PITA-Sb pairing exhibited notable changes in most primary metabolite groups, while other EM pairings exhibited more scattered responses (Fig. 2A). Both increases and decreases in carbohydrates, amino acids, and organic acids by mycorrhizas were observed (Fig. 2A).

**Fig. 2.**
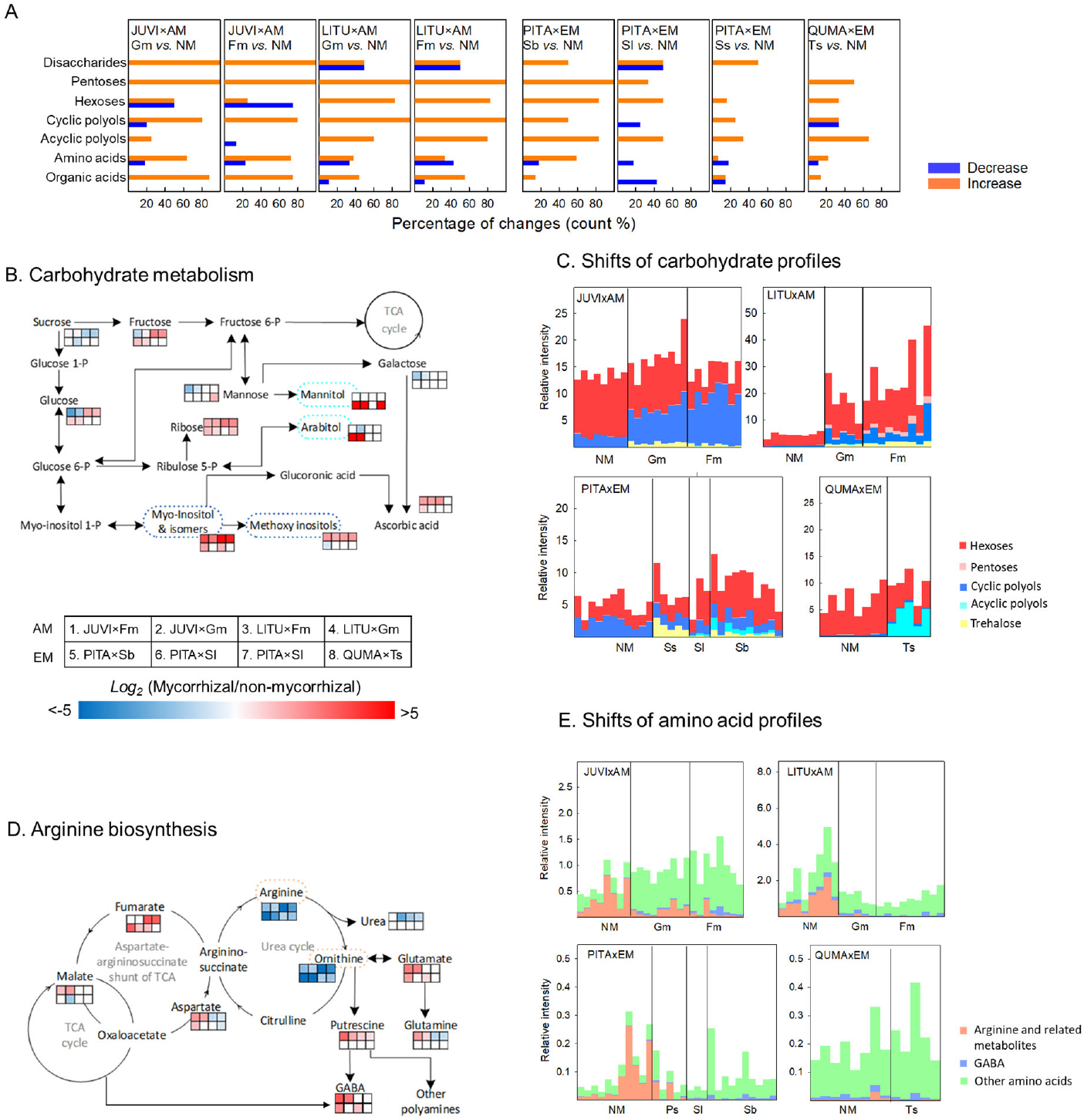
Effects of mycorrhizal associations on root primary metabolism in eight plant-fungus combinations across AM and EM lifestyles. **A.** Percentage of changes (two-tailed Student’s *t*-test, *P* < 0.05) in primary metabolites. **B.** Sugar metabolism simplified from KEGG primary pathways. **C.** Shifts of carbohydrate profiles were shown by the summed relative intensity (normalized to the internal standard) of carbohydrates and polyols across AM and EM lifestyles. **D.** Arginine biosynthesis pathway simplified from KEGG amino acid pathways. **E.** Changes of amino acid profiles were shown by the summed relatively intensity (normalized to the internal standard) across AM and EM lifestyles. In **B.** and **D.**, the compounds with significant differences (*P* < 0.05) between mycorrhizal and non-mycorrhizal roots are shown in a gradient from red to blue colors, with blank indicating non-significant effects or undetected compounds.

Mycorrhizas frequently affected root carbohydrate profiles (Fig. 2BC, and S2). The pathway analysis detected significant alterations on several sugar metabolism pathways, with galactose metabolism being the most sensitive, exhibiting a significant change in six out of eight plant-fungal combinations across AM and EM (Table S1). Particularly, the abundance of carbohydrates such as sucrose, glucose, fructose, and mannose were frequently affected by symbiosis, but their responses were highly variable across plant-fungal combination (Fig. 2B, S1). For example, the responses of the two hexoses that are readily transportable from roots to mycorrhizal fungi (*i.e*., glucose and fructose) differed among plant species: AM decreased glucose and fructose in JUVI roots but increased these compounds in LITU roots; EM increased glucose and fructose in PITA roots but they were unaffected in QUMA roots (Fig. 2B, S2).

In contrast to the variable responses of hexose currency, comparing metabolite alterations across plant-fungus combinations reveals a similar trend in another major C pool of roots, *i.e*., polyols and their derivatives, within one mycorrhizal lifestyle (Fig. 2BC, S2). Consistent with hypothesis (1), AM and EM lifestyles tended to affect polyols in different manners. The alterations of root carbohydrates between non-mycorrhizal and AM-colonized roots featured an accumulation of cyclic polyols, *i.e*., inositols and their methoxy derivatives such as ononitols and pinitols (Fig. 2BC, S2). This accumulation occurred in JUVI and LITU with AM lifestyles irrespective of phylogenetic history of plant hosts (*i.e*., gymnosperm *vs*. angiosperm) or fungal identities (*i.e*., Fm or Gm,). By contrast, EM-colonized roots did not exhibit similar increases in cyclic polyols but substantially accumulated a set of new compounds that were undetected in non-mycorrhizal roots, including acyclic polyols and trehalose (Fig. 2BC, and S2). The large increases of acyclic polyols in EM-colonized roots were driven by the accumulation of mannitols and arabitols in PITA and mannitols in QUMA roots (Fig. S2CD), where these compounds contributed >96% of the increase in acyclic polyols. The only pairing of EM association (PITA × Ss) that did not accumulate acyclic polyols exhibited a large increase of trehalose (Fig. 2C).

Variance partitioning analysis showed that the observed changes in root carbohydrates were more related to mycorrhizal colonization than plant growth and tissue N (Fig. S3). Overall, mycorrhizal colonization explained 22 – 64% of the variation in the profiles of polyols, with 9-21 % of the variation explained by mycorrhizas independently (*P* < 0.063), whereas growth and tissue N generally did not show significant and unique effects (*P* ≥ 0.308, Fig. S3). To understand if the changes of polyols by mycorrhizas occurred in a localized or systemic manner, we also investigated the abundance of polyols in the corresponding leaves in each plant-fungus combination. We observed increases of cyclic polyols in leaves of AM-inoculated JUVI seedlings, although to a considerably lesser degree when compared to their belowground counterparts (Fig. 2C, S4A). Acyclic polyols such as mannitols and arabitols were undetected in the leaves of EM-inoculated seedlings (Fig. S4A). These results indicate that the observed accumulation of polyols in colonized roots was largely driven by mycorrhizas and the accumulation of acyclic polyols by EM colonization was localized in roots.

Amino acids and organic acids also varied between non-mycorrhizal and mycorrhiza-inoculated roots (Fig. 2DE, and S2). Although the responses of amino acids and organic acids were generally mixed, we observed a common trend where the abundance of arginine and its catabolism products such as ornithine and urea were consistently decreased by mycorrhizas across most plant-fungus combinations irrespective of mycorrhizal lifestyles, leading to a characteristic shift of amino acid profiles associated with mycorrhizas (Fig. 2DE, and S2). Pathway analysis also showed that arginine biosynthesis was highly sensitive to mycorrhizal symbiosis and was altered by symbiosis in seven out of eight plant-fungal combinations (Table S1). Particularly, metabolites involved in the urea cycle (*i.e*., arginine, ornithine, and urea) were significantly decreased in most plant-fungal combinations (*P* < 0.023, Fig 2DE), while the metabolites from their neighboring pathways, *e.g*. the biosynthesis of polyamines and γ-aminobutyric acid (GABA, a stress-related non-protein amino acid), were generally increased, indicating C flow away from the urea cycle (Fig. 2DE). Variance partitioning showed that the decrease in arginine & related metabolites were predominantly driven by tissue N, which independently explained 18–31% of the variation (Fig. S3). The decrease in arginine and its catabolism products also occurred in the leaves of mycorrhiza-inoculated seedings (Fig. S4B), indicating a systemic response.

### Alterations of specialized metabolism

To understand the broad response of specialized metabolism, we first combined LC-MS data with *in silico* chemical taxonomy assignment (*see* Methods). This procedure annotated a large proportion of the detected mass features (at class level: 71 % to 88 %, *e.g*., flavonoids; at level 5: 47 - 62%, *e.g*., triterpene glycosides) and revealed a similar distribution of root metabolome at high levels of chemical classes across plant species (Fig. 3A). For example, at the superclass level, the root metabolome from all four plant species was dominated by organic acids & derivatives, organic oxygen compounds, lipids & like, and phenylpropanoids & polyketides. Flavonoids were a major group of phenylpropanoids & polyketides across plant species, whereas prenol lipids and fatty acyls comprised the majority of lipids & like. This broad similarity in root chemistry provides opportunities for understanding the potential commonality vs. specificity of metabolite alterations. Root metabolomes also exhibited plant species-specific chemical characteristic, especially at finer levels (Fig. 3A), reflecting interspecific chemical plasticity. For example, phenylpropanoids were predominantly dominated by flavonoids in PITA roots, LITU roots produced similar numbers of flavonoids and cinnamic acids & derivatives, while QUMA roots were abundant in hydrolysable tannins (Fig. 3A).

**Fig. 3.**
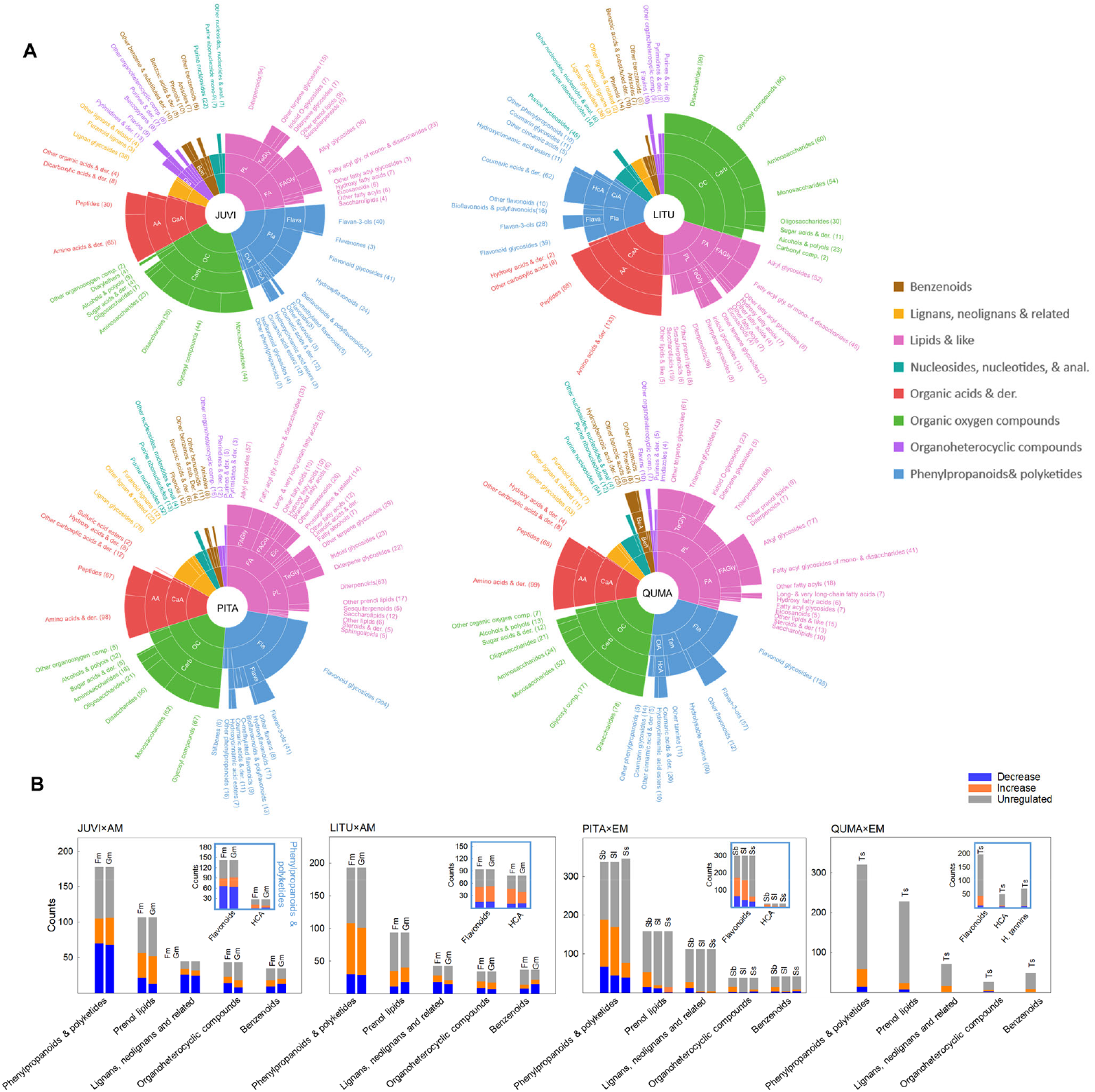
Effects of mycorrhizal associations on root specialized metabolites in eight plant-fungus combinations across AM and EM lifestyles. **A.** The extracted mass features were assigned and grouped into four chemical taxonomy levels (from superclass, class, subclass, to level 5 outward, see http://classyfire.wishartlab.com/ for details) according to a computing-assisted procedure CANOPUS (*see* Method). AA: Amino acids, peptides, & analogues; BeA: benzenoid acids & derivatives (der.).; Ben: benzene and substituted der.; CaA: carboxylic acids & der.; Carb: carbohydrates & conjugates; CiA: cinnamic acids & der..; Dia: diazines; Eic: eicosanoids; FA: fatty acyls; FACoj: fatty acids & conjugates; FAGly: fatty acyl glycosides (gly.); Fla: flavonoids; Flava: flavans; HcA: hydroxycinnamic acids & der.; OC: organooxygen compounds (comp.); PL: prenol lipids; Tan: tannins; TeGly: terpene gly. Groups with features <2 were combined for better comprehensibility. **B.** The counts of increases, decreases (two-tailed Student’s *t*-test, *P* < 0.05), and unregulated mass features in response to mycorrhizal symbiosis are shown for major specialized metabolite groups. The insets show the changes (*P* < 0.05) of subgroups within the chemical class of phenylpropanoids & polyketides. HCA: hydrocinnamate acids and derivatives; H. tannins: hydrolysable tannins.

Most of the *in silico* assigned specialized metabolite groups exhibited significant changes in response to mycorrhizas (Fig. 3B). AM tended to elicit higher numbers and greater proportions of changes than EM: extensive increase and decrease within each group were observed in both AM host species (Fig. 3B). Among metabolite groups involved in specialized metabolism, *in silico* assigned phenylpropanoids exhibited the highest numbers of mycorrhiza-associated changes (Fig. 3B). The responsiveness of phenylpropanoids was relatively consistent across plant-fungus combinations: although EM roots tended to be less responsive than AM roots (*e.g*., regarding prenol lipids, lignans, and benzenoids), the EM fungi Sb and Sl still elicited a high number (169 to 189) and a large proportion (50.0 to 55.9%) of responses within phenylpropanoids in PITA roots (Fig. 3B). By contrast, although prenol lipids showed a relatively high response to AM (particularly, diterpenoids in JUVI and terpene glycosides in LITU), EM elicited considerably lower proportions of responses in this group of compounds (Fig. 3B). The high response of phenylpropanoids to symbiosis were predominantly driven by the changes in the mass features assigned as flavonoids, which accounted for 86.4%, 42.5%, 90.8%, 70.1% of the observed responses of phenylpropanoids in JUVI, LITU, PITA, and QUMA, respectively (Fig. 3B). Accordingly, we focused on flavonoids & related metabolites because: **1)** this chemical family was highly responsive to mycorrhizas; **2)** flavonoids are almost exclusively synthesized by plants. Our survey across the genome sequences of the studied fungal species/genus (*i.e*., Gm, Sb, Sl, and Ts) suggests that none of these fungi possesses the essential enzyme kit (ammonia-lyase, chalcone synthase, and chalcone isomerase, Winkel 2006) required in the flavonoid biosynthesis (*see* the database: MycoCosm https://mycocosm.jgi.doe.gov/mycocosm). Although we did not separate plant from fungal metabolites, the exclusiveness of flavonoids allows for elucidating modulation of plant specialized metabolites without confounding changes by fungal metabolism. In addition, **3)** the core pathway of flavonoid metabolism is largely shared across the plant kingdom (Wen et al., 2020), allowing for comparing the responses of these metabolites across various plant-fungus systems.

We identified 66, 48, 67, 80 specialized metabolites for JUVI, LITU, PITA, and QUMA respectively based on accurate mass and MS^n^ spectra, which broadly cover the phenylpropanoid, flavonoid, and the adjacent metabolism pathways (Fig. 4). The details of annotation are shown in Table S2, and the list of putative annotates for each plant-fungus combination and their respective responses in Table S3. Pathway analysis based on annotates showed that flavonoid biosynthesis and flavone and flavonol biosynthesis were frequently affected by mycorrhizas across plant-fungal combinations (Table S1). Comparison of specialized metabolic responses across plant-fungus combinations revealed consorted, yet differential responses between different flavonoid subgroups, a pattern in line with hypothesis (2). The members of a major flavonoid subgroup, flavonols (*e.g*., quercetins, kaempferols, myricetins) and their derivatives exhibited mixed, largely decreasing responses to mycorrhizas (Fig. 4AB, Table S3). By contrast, we observed a generally consistent increase in the members of another major flavonoid subgroup corresponding to flavan-3-ols (*e.g*., catechins, gallocatechins) and their oligomers (*e.g*., procyanidins and prodelphinidins, Fig. 4AB, Table S3), leading to a pronounced increase in the summed peak intensities of this group in seven out of eight plant-fungal combinations (*P* < 0.001, Fig. 4B). The consistent accumulation of flavan-3-ols across plant-fungal combinations along with the general decrease of other flavonoids resulted in a characteristic shift of flavonoid profiles by mycorrhizas in most plant-fungal combinations irrespective of mycorrhizal lifestyles (Fig. 4B). The only exception is the PITA-Sl pairing, where the colonization rate was the lowest among plant-fungal combinations (Table 1). Mapping the annotated compounds to the flavonoid biosynthesis pathway showed that forming mycorrhizas tended to drive C flow towards later steps of the flavonoid biosynthesis pathway, as flavan-3-ols are synthesized further downstream than flavonols (Fig. 4A). Variance partitioning showed that mycorrhizal colonization was the most important predictor for the changes of flavan-3-ols, explaining 56% - 86% of the variation, with 9% - 81% of the variation explained by mycorrhizal colonization independently (Fig. S3). The accumulation of flavan-3-ols in mycorrhiza-colonized roots did not mirror in their corresponding leaves (Fig. S4). Instead, the presence of belowground symbiosis did not affect or slightly decreased the summed peak intensities of flavan-3-ols in leaves.

**Fig. 4.**
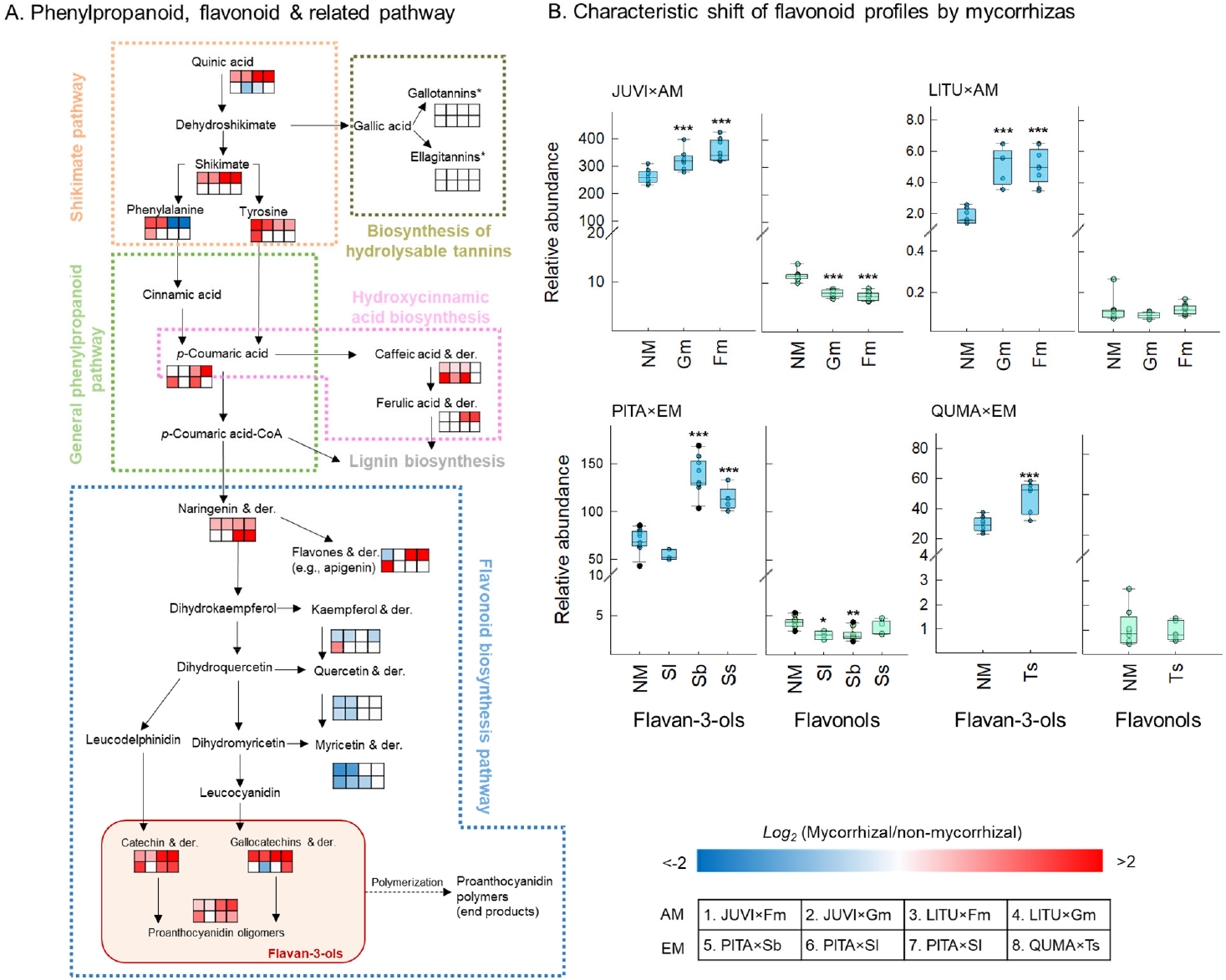
Characteristic shifts of flavonoid profiles in response to mycorrhizal associations in eight plant-fungus combinations across AM and EM lifestyles. **A.** Flavonoid biosynthesis simplified from KEGG pathways. Compounds with significant differences (two-tailed Student’s *t*-test, *P* < 0.05) between mycorrhizal and non-mycorrhizal (NM) roots are shown in a gradient from red to blue colors, with blank indicating non-significant effects. *We note that gallotannins and ellagitannins were only present in QUMA roots and did not show significant responses to mycorrhizas. **B.** Characteristic shifts of flavonoid profiles in response to mycorrhizal symbiosis featured by the differential changes in the summed intensity (normalized to internal standards) between the subgroups of flavan-3-ols and flavonols. See the list of annotated flavan-3-ols and flavonols in Table S3. * (mycorrhizal *vs*. NM): *P* < 0.05; **; *P* < 0.01.***; *P* < 0.001.

Because the number of annotates is limited compared to that of the extracted mass features, we further tested if this characteristic shift of flavonoid profiles associated with mycorrhizas is robust to a broader range of *in silico*-assigned flavonoids. Similarly, the mass features assigned *in silico* as flavonols and their derivatives exhibited mixed, largely decreasing responses to mycorrihzas; by contrast, we observed generally consistent increases in the mass features assigned as flavan-3-ols (Fig. S5AB). The summed peak intensities of mass features assigned as flavan-3-ols increased by mycorrhizas in seven out of eight plant-fungus combinations, whereas that of the mass features assigned as other flavonoids generally decreased (Fig. S5C). These patterns of *in silico* assigned mass features further support the shift of flavonoid profiles (*i.e*., accumulation of flavan-3-ols) as a characteristic alteration of specialized metabolites associated with mycorrihzas irrespective of mycorrhizal lifestyles.

## Discussion

Characterizing the chemical landscapes associated with plant-mycorrhizal interactions using comprehensive metabolomics, our data showed that the chemical responses of root metabolome to mycorrhizas were highly complex, involving decreases and increases in a large number of metabolites spanning major chemical classes (Fig. 1–4). However, using a multi-species approach, common *vs*. lifestyle-specific metabolite alterations across various plant-fungus combinations have emerged. Although mycorrhiza-associated metabolite responses in plants are often considered highly specific to plant and/or fungal species (Schweiger et al., 2014), our data revealed for the first time, to our knowledge, that part of root metabolite alterations by mycorrhizas were relatively common across plant-mycorrhizal systems; this commonality is particularly striking as it persisted irrespective of the phylogenetic relationships among host plants and appeared robust to various mycorrhiza-mediated changes on plant growth and N status.

Similar to mycorrhizas-associated common traits at the genetic level, the common metabolite alterations by mycorrhizas within one or across mycorrhizal lifestyles may also reflect selection pressures and biological strategies critical for mycorrhization in general. For example, carbohydrate partitioning between plant hosts and fungal partners is fundamental for successful symbiosis, as carbohydrates are the currency that plants use for exchange with fungi. Mutualistic/balanced plant-mycorrhizal associations would require well-regulated C partitioning between plants and fungi (Brundrett, 2002). Such a general requirement may result in convergent alternations in root carbohydrate profiles. However, the response of the immediate exchange currency (*i.e*., glucose and fructose) were highly variable across plant-fungus combinations (Fig. 2, S2), reflecting the dynamic nature of these carbohydrates as they are readily transportable. On the other hand, our data revealed that polyols, a major C pool in root tissues, consistently accumulated in mycorrhiza-colonized roots. The two AM hosts, although phylogenetically and chemically distinct from each other (Fig. S1), accumulated cyclic polyols (inositols & derivatives) that typically have a plant origin (*see* the BinBase database, Lai et al., 2018; also *see* Fig. 2, S3 that show the presence of inositols in uninoculated roots and leaves). Accumulating large amounts of cyclic polyols that are unavailable for symbionts may help plants to mitigate the hexose gradient at the root-fungal interface and thus regulate carbohydrate efflux to AM (Nehls et al., 2007; Nehls and Bodendiek, 2012). In line with our observation, at cellular-level, inositols are among the most responsive carbohydrates to AM: arbuscule-containing cells accumulated inositols while hexoses were generally unaffected (Gaude et al., 2015). A recent study also showed that accumulation of inositols was pronounced in mutualistic AM associations but was lacking when AM fungi became parasitic (Kaur et al., in revision). These and our observation together suggest that accumulation of inositols may be a common C partitioning strategy in AM-associated roots and open intriguing questions on the potential role of such a frequent accumulation (*e.g*., a possible avenue for plants to regulate C exchange with fungi). By contrast, consistent with hypothesis (1), EM-associated roots accumulated a different set of polyols (acyclic polyols such as mannitols and arabitols, Fig 2, S2) that often have a fungal origin and thus may have different biological consequences. Mannitols/arabitols are known to be produced in large quantities by EM fungi (Nehls and Bodendiek, 2012). The EM host PITA does not normally produce mannitols and was trans-genetically introduced with genes involved in synthesizing mannitols to enhance stress tolerance (*e.g*., Tang et al., 2005). In addition, mannitols/arabitols did not occur in uninoculated roots and leaves (Fig. S4). Therefore, their frequent accumulation in EM-inoculated roots was more likely driven by fungal metabolism. In contrast to the accumulation of inositols in AM roots that may help plants regulate C efflux, quickly converting hexoses to acyclic polyols may instead help EM fungi maintain a strong sink for carbohydrates (Lewis and Harley, 1965; Nehls et al., 2007; Nehls and Bodendiek, 2012). Interestingly, accumulation of mannitols/arabitols has also been observed in fungal pathogens as important for pathogenicity, with proposed functions such as carbohydrate reserves and quenchers for plant ROS responses (Patel and Williamson, 2016; Hooshmand et al., 2020). This similarity between EM and pathogens implicates an evolutionary scenario where the accumulation of these acyclic polyols could be crucial for the establishment of fungal colonization in plant tissues. Taken together, the contrasting alterations in carbohydrate profiles between AM and EM lifestyles reflect their unique C partitioning strategies for sustaining symbiosis and may provide potential mechanisms driving the divergent C utilizations between these mycorrhizal lifestyles.

Alterations of specialized metabolites that regulate plant defense to biotic/abiotic stresses also represent an essential step for mycorrhization. To facilitate symbiosis, plant defense responses were often suppressed as a result of decreased expression of defense-related genes (Zamioudis and Pieterse, 2012). On the other hand, plants need to delimit the undesirable spread of mycorrhizal fungi by modulating tissue chemistry; this modulation is more likely through altering the specialized metabolites than strengthening physical barriers, as symbiosis often precedes significant lignin/suberin deposition in the endodermis (Brundrett, 2002). Therefore, the general challenges of mycorrhization (facilitation *vs*. delimitation) could be translated into complex alterations in specialized metabolites. Our data showed that a large specialized metabolite group, flavonoids, are highly responsive to mycorrhizas (Fig. 3, 4). The core steps of flavonoid biosynthesis pathway are largely conserved (Fernie, 2019), which may allow for convergent responses to mycorrhizas across plant species. Indeed, comparing across multiple plant-fungus combinations revealed a characteristic shift in flavonoid profiles that occurred frequently in mycorrhizal colonized-roots irrespective of lifestyles (Fig. 4). In line with hypothesis (2), this shift featured a general decrease in flavonols but an increase in flavan-3-ols, which may be interpreted in the context that plants both facilitate and delimitate fungal growth during mycorrhizal interactions. Flavonols and their derivatives exhibited a variable, generally decreasing trend in colonized roots (Fig. 4, S5, Table S3). The mixed responses of flavonols suggest that this flavonoid family may have versatile functions. Indeed, while flavanols could have antifungal activity (*e.g*., glycosylated quercetin, Sudheeran et al., 2020) and thus are at odds with mycorrhization, certain flavonols can instead stimulate mycorrhizal growth and positively relate to colonization (Lagrange et al., 2001). In contrast, the family of flavan-3-ols (*e.g*., catechins, gallocatechins, and their derivatives) showed a consistent accumulation across various plant-mycorrhizal combination irrespective of mycorrhizal lifestyles. These increases were unique to roots and did not occur in leaves (Fig. 4, S4), indicating an essential regulatory role of this flavonoid subgroup in root-mycorrhizal interactions. Flavan-3-ols are among the most potent flavonoids to defend against fungal penetration (Friedman, 2007; Ullah et al., 2019). Catechins (often a dominant type of flavan-3-ols in root tissues) were observed to accumulate in large quantities in the inner part of root endodermis in *Larix decidua* Mill. (Weiss et al., 1997, 1999) and cotton seedlings (Mace & Howell, 1974), forming critical barriers against fungal inward growth to root vascular tissues. Hence, the general accumulation of flavan-3-ols in AM- and EM-colonized roots observed in this study may represent a common defense investment of plant hosts to prevent undesirable fungal penetration. In addition, the protective role of flavan-3-ols may be multi-level. Many members of this group are efficient ROS scavengers that can trap an additional peroxyl radical compared to other flavonoids (Kondo et al., 2000). Their antioxidant properties may protect plant tissues from the fungal-induced ROS defense responses. Imaging studies that examine the localization of flavan-3-ols in colonized roots could further elucidate their protective roles in root-mycorrhizal interactions.

The common metabolite responses observed in roots were largely localized and did not mirror in leaves (Fig. S4). This may explain the lack of commonality across plant species in leaf metabolic responses to *Rhizophagus irregularis* (Schweiger et al., 2014). Leaves are not the immediate site of mycorrhization or directly respond to the unique challenges (*e.g*., delimiting the undesirable fungal spread) that may shape root chemistry in a similar way and lead to convergent metabolite alterations. Therefore, the chemical composition of leaves may not be as tightly connected to mycorrhizas as those in roots (Xia et al., 2021). However, our data supported that symbiosis could induce systemic changes that indirectly affect leaf chemistry. For example, the decrease in arginine & related metabolites occurred in both roots and leaves and was more related to tissue N than mycorrhizal colonization (Fig. 2, S4), suggesting that these changes in leaves were driven by the mycorrhiza-induced systemic changes in plant N utilization. Our data demonstrated that mycorrhizas have different relationships to root and leaf chemistry and thus the transferability of the results in the metabolic responses to mycorrhizas from roots to leaves (and *vice versa*) is limited.

In conclusion, we demonstrated common vs. lifestyle-specific traits of metabolite alterations across various plant-mycorrhizal combinations. Although many metabolites (*e.g*., hexose, flavonols) showed mixed responses that were specific to plant species and/or fungal identities, a subset of highly responsive compounds (*e.g*., polyols and flavan-3-ols) exhibited common responses to mycorrhizas that were unique to or shared across AM and EM lifestyles, highlighting their potentially critical regulatory role in root-mycorrhizal interactions. These common patterns appear robust to the phylogenetic diversity of the host plants, and thus may be widespread in land plants. This study offers future research venues to elucidate the finer roles of these common traits of mycorrhiza-associated metabolite alterations and thus help to eventually decipher the nature of this widespread plant-fungus partnership.

## Materials and Methods

### Greenhouse experiments

We chose phylogenetically-diverse host trees to maximize our understanding of mycorrhiza-mediated chemical changes. Specifically, we included angiosperms and gymnosperms that are host plants for both mycorrhizal types to help reconcile the underrepresentation of AM-associated conifers in literature and to compare the potential shaping role of phylogenetic relationship *vs*. mycorrhizal lifestyle on metabolite alterations. We inoculated each of these host plants with multiple fungal species to assess how responses may differ by specific plant-fungal pairings. All plants were grown in the Plant Growth Facilities greenhouse at the University of Minnesota (UMN), St. Paul, MN, USA. The seeds of AM-associating *Juniperus virginiana* (JUVI) and *Liriodendron tulipifera* (LITU), and the EM-associating *Pinus taeda* (PITA) were purchased from Sheffield’s Seed Company (Locke, NY, United States). The EM-associating *Quercus macrocarpa* (QUMA) was collected on the UMN campus grounds. The AM fungal inoculums *Gigaspora margarita* (Gm) and *Funneliformis mosseae* (Fm) were purchased from MycoBloom (MycoBloom LLC, Lawrence, KS, USA). The EM fungal inoculums included *Suillus brevipes* (Sb), *Suillus luteus* (Sl), *Thelephora terrestris* (Tt), *Laccaria sp*. (Ls), and *Tuber sp*. (Ts). Spores for the EM inoculums were prepared from field-collected sporocarps with spore slurries made following Mujic *et al*. (2016), with a small section of gleba was subjected to DNA extraction for species identification. The identity of each EM fungal species was further confirmed by DNA analysis for the colonized root tissues after harvest (details are shown in the next subsection).

Seeds for all hosts but QUMA were placed on a cloth bed, covered with cloth, dipped in 8% H_2_O_2_ for 30 minutes, and soaked with H_2_O for 24 h. The bedded seeds were incubated at 4 °C for stratification before sowing (60, 80, 180 days for PITA, JUVI, and LITU, respectively). For QUMA, germinated acorns with radicals not penetrating the soil were collected in September 2018 and rinsed with a 10% bleach solution before sowing. The soil (50:50 mix of commercial sand and soil from the Cedar Creek Ecosystem Science Reserve, East Bethel, MN) was autoclaved and added to 70%-ethanol sterilized plastic trays, with 2.5 L soil for each tray. Inoculums were added at a rate of 10 % v/v. The two AM hosts were each inoculated with Gm and Fm; the two EM hosts were each inoculated with Ts, and Ls, while PITA was additionally inoculated with Tt, Sb and Sl. All mycorrhizal species were added as separate, single-species inoculums in separate trays. Each plant-fungal combination was represented in three separate trays along with three trays for non-inoculated seedlings for each host species. Turf Partners fertilizer was applied at the lowest recommended concentration monthly to each tray. Seedlings of JUVI and LITU were harvested after ca. seven months of growing, while PITA and QUMA were harvested at 9 and 11 months.

During harvest, plant height (the length from the soil surface to apical meristem) for each seedling was recorded to estimate plant growth. Leaves and absorptive roots were collected from three to five individuals in each tray, and roots were separated into two subsamples for separate chemical and mycorrhizal analysis. The samples used for chemical analysis were immediately flash-frozen in liquid nitrogen and stored at −80 °C until shipping to Clemson University with dry ice. The samples to be scored for colonization were placed in 60% ethanol until analysis. The remaining leaf and root material in each tray were collected and dried for 48 hours at 35 °C. We collected root materials for DNA analysis to confirm fungal identity.

### Mycorrhizal colonization and fungal identity

Samples stored in ethanol were washed in dH_2_O and placed in glass vials. AM-inoculated roots were cleared by adding 10% KOH and autoclaving for a 60-min liquid cycle, which were then rinsed with dH_2_O and acidified using 2.5 % HCl for 10 min. Once acidified, Trypan Blue solution was added to each vial and set to autoclave for a 60-min liquid cycle. Colonization of each seedling was quantified using gridline-intersect method (Giovannetti and Mosse, 1980). For EM colonization, all root tips in each sample were scored under a dissecting microscope. In the Ls treatments for PITA and QUMA, and Ts for PITA, only a trace amount of colonization on a few seedlings was present, so they were excluded in chemical analyses. The colonization of PITA roots by Sl was also relatively low (~5%). However, we included Sl-associated roots for chemical analysis as this low colonization resulted in an improved growth (Table 1) and a shift in root chemistry (Fig. 1). Prior studies have also shown that low EM colonization could cause improved plant growth and significant readjustment of plant metabolites (Szuba et al., 2020).

Molecular identification of EM fungal identities was done by first extracting DNA using the REDEextract-N-Amp protocol (Sigma–Aldrich, St. Louis, MO, USA). PCR using the fungal-specific primer pair ITS1f-IT4 was conducted using the thermocycling conditions detailed in Kennedy *et al*. (2020). PCR products were cleaned with Exo-SAP IT (USB Corp., Cleveland, Ohio, USA) and sequenced in forward direction with the ITS1F primer at the UMN Genomics Center. Sequences were queried against the NCBI database to ascertain fungal identity. The Sb, Sl, and Ts treatments matched closely (>99% similarity) to the species used in the inoculums, but the EM root tips in the Tt treatment matched closely to an Sordarialean EM fungus previously encountered in artic EM vegetation surveys (Brevik et al., 2010). Since colonization by this fungus was consistently high (~74.6% by average, Table 1), we included this treatment (coded as Ss) in chemical analyses.

### Chemical analysis and data processing

To correctly link the chemical changes to colonization by each fungal inoculum, seedlings were first examined for mycorrhizal colonization and fungal identity. Only individual plants exhibiting mycorrhizal colonization by the identified fungal species were analyzed for chemistry. Accordingly, seedlings from the three trays for each of the JUVI-Gm, JUVI-Fm, LITU-Fm, PITA-Sb pairs were included in chemical analysis (biological replicates n = 9-10, Table 1). Mycorrhizal colonization only occurred in two trays for the LITU-Gm (n = 5) and QUMA-Ts pairs (n = 6), and in one tray for the PITA-Ss pair (n = 5) and the PITA-Sl pair (n = 3). We did not observe mycorrhizal colonization in non-mycorrhizal controls (n=8-11). Chemical analysis was performed for roots and leaves, except for LITU seedlings where only roots were analyzed as we did not collect sufficient leaf material for chemical analysis.

For chemical analysis, 20 mg of the freeze-dried and cryohomogenized samples were extracted with 500 μl methanol under sonication for 1 h with ice. 50 μl methanol extractants from each sample were spiked with a 50-μl internal standard (1 μg ml^−1^ resveratrol, 99 atom % ^13^C6, Sigma-Aldrich, St. Louis, MO, USA), and analyzed using an Ultimate 3000 HPLC (Thermo Scientific, Waltham, MA, USA) by an Acquity UPLC HSS T3 column (150 × 2.1 mm, 1.8 μm; Waters Corp., Milford, MA, USA), using a gradient elution of 0.1% formic acid and acetonitrile. The compounds were then analyzed using an Orbitrap Fusion Tribrid Mass Spectrometer (Thermo Scientific Inc., Waltham, MA, USA) equipped with an electrospray ion source (UHPLC-Orbitrap-MS/MS). The mass spectrometer was operated in negative ionization mode with a data-dependent fragmentation (MS2) using a HCD dissociation mode (Bowers et al., 2018). Details of UHPLC-Orbitrap-MS/MS analysis are shown in Method S1.

We processed the LC-MS files with Compound Discoverer (ver 3.2, Thermo Fisher Scientific, Waltham, MA, USA) for feature extraction, chromatogram deconvolution, isotope grouping, alignment, and putative compound annotation. The workflow extracted 6920 to 10834 individual mass features for each plant species. Putative molecular formulas were estimated from most of the features (> 98%) using 5 ppm as a threshold. The elemental ratios derived from these formulas were then used to construct *van* Krevelen diagrams based on the constraints of O:C and H:C ratios (Brockman et al., 2018; Rivas-Ubach et al., 2018). Putative annotation was based on accurate mass (< 5ppm) and MS^n^ fragmentation according to the mzCloud library and literature. The details of annotation and mass features of each annotated compound were shown in Table S2. We also processed the LC-MS files in MZmine 2.53 for computing-assisted class-level annotation. This workflow extracted 1118 (JUVI) to 2293 (QUMA) mass features which were summarized in .mgf format and a feature quantification table and uploaded to the GNPS for the feature-based molecular networking (FBMN) workflow, as an attempt to structure the complex metabolome based on spectral similarity (Nothias *et al*., 2020). The FBMN workflow was performed with an ion tolerance of 0.002 Da, a cosine score of 0.7, and a maximum shift between precursors of 500 Da. The .mgf file for SIRIUS platform (4.8.2) exported from MZmine was used to predict compound classes with the *in silico* CANOPUS workflow (Dührkop *et al*., 2021) for unannotated chemical features, using automated structure-based chemical taxonomy (ClassyFire, Feunang et al., 2016) to multiple levels (Superclass, Class, Subclass, and Level 5, *see* http://classyfire.wishartlab.com/ for details of chemical taxonomy). The molecular networking from FBMN and chemical classes assigned by CANOPUS were combined in Cytoscape (3.8.2) for visualization.

We performed a separate metabolite profiling using GC-MS analysis to better characterize polar metabolites that cover most of the primary metabolites (*e.g*., soluble carbohydrates, amino acids, Suseela et al., 2015). 200 μl methanol extract was phase-separated in a chloroform-water-methanol system. 20 μl of the water-methanol phase from each sample and a gradient of standard solutions containing 38 authentic compounds (Table S2) were each spiked with a 10-μl internal standard (50 μg ml^−1^ myristic acid + 20 μg ml^−1^ ribitol) and dried under vacuum. The dried samples were methoximated with 20 μl Methoxyamine-HCl (40 mg ml^−1^ in pyridine) and derivatized by 80 μl MSTFA + 1% TMCS with 1 μg ml^−1^ alkanes (C10-C34) before being analyzed on a gas chromatography-quadrupole time of flight mass spectrometry (Agilent 7250 GC-QTOF-MS, Agilent Technologies Inc., USA). The GC-MS files were processed with MS-DIAL (4.48) for peak detection, deconvolution, compound identification, and alignment (within one host species). The annotation of compounds was based on comparing the kovats retention index built on n-alkanes standards (C10-C34) and the mass spectra with authentic standards, a database (MS-DIAL metabolomics MSP), and/or literature. Details of annotation were shown in Table S2. Carbon and N concentrations of plant roots and leaves were determined by a Carlo Erba Elemental Analyzer at Duke University.

### Data analysis

Data analysis was performed with R software (version 4.0.3) unless otherwise stated. For the principal coordinate analysis (PCoA), the dissimilarity index matrix (Euclidean) was first constructed using the function *vegdist* from the vegan package after data standardization. Then, the function *pcoa* from the package ape was used to compute the principal coordinate decomposition of this distance matrix. To test the significance of clustering pattern by treatments, we performed a permutational multivariate analysis of variance (PERMANOVA) based on all significant axes using the function *adonis* from the package *vegan*. The significance of differences in colonization rates, plant heights, and nutrient concentrations were tested with general linear models using the function *lm* from the R-core package followed by *post hoc* comparisons. However, when the non-normality and homogeneity of variance assumptions were not met, Kruskal-Wallis tests were performed to examine the main effect, followed by individual pairwise Mann Whitney U tests for multiple comparison. The pairwise comparison (mycorrhizal *vs*. non-mycorrhizal) of chemical features was examined by a two-tailed Student’s *t*-test on the *log*-transformed peak intensities. The variance partitioning that compared the potential drivers of the metabolite responses followed the Legendre method (Legendre & Legendre, 2012) using the function *varpart* from package vegan. Data were *log*-transformed and standardized prior to multivariate analyses. The pathway (KEGG) analysis was performed with PaintOmics 3 (v0.4.5, García-Alcalde *et al*., 2011) and Fisher exact tests were used to test the significance of mycorrhizal effects on pathways.

## Acknowledgments

This research is supported by funding from the grants (DEB 1754679) from the U.S. National Science Foundation.

## Supplementary files

**Fig. S1.**
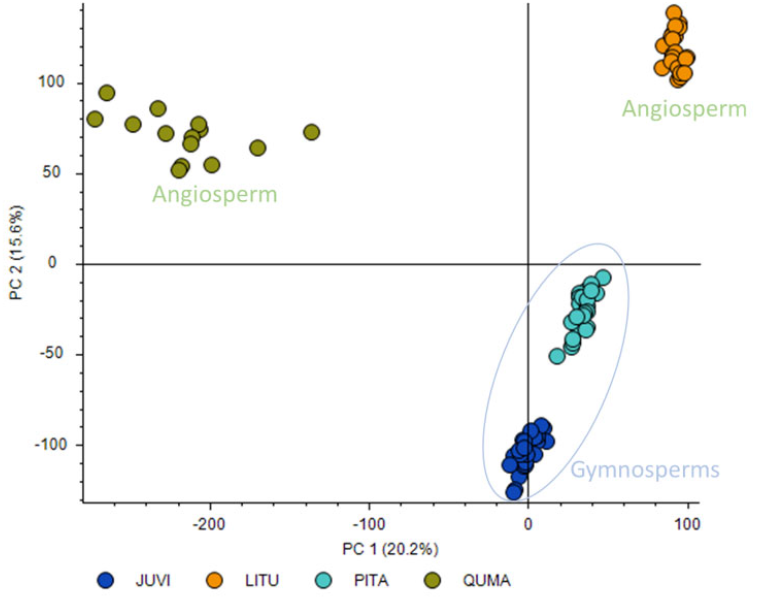
Distinct root chemical phenotypes of the four plant species revealed by principal coordinates analysis. The two gymnosperms *Juniperus virginiana* (JUVI) and *Pinus taeda* (PITA) are located closer to each other while farther away from the two angiosperms *Liriodendron tulipifera* (LITU) and *Quercus macrocarpa* (QUMA) in the chemical space spanned by the first two latent components.

**Fig. S2.**
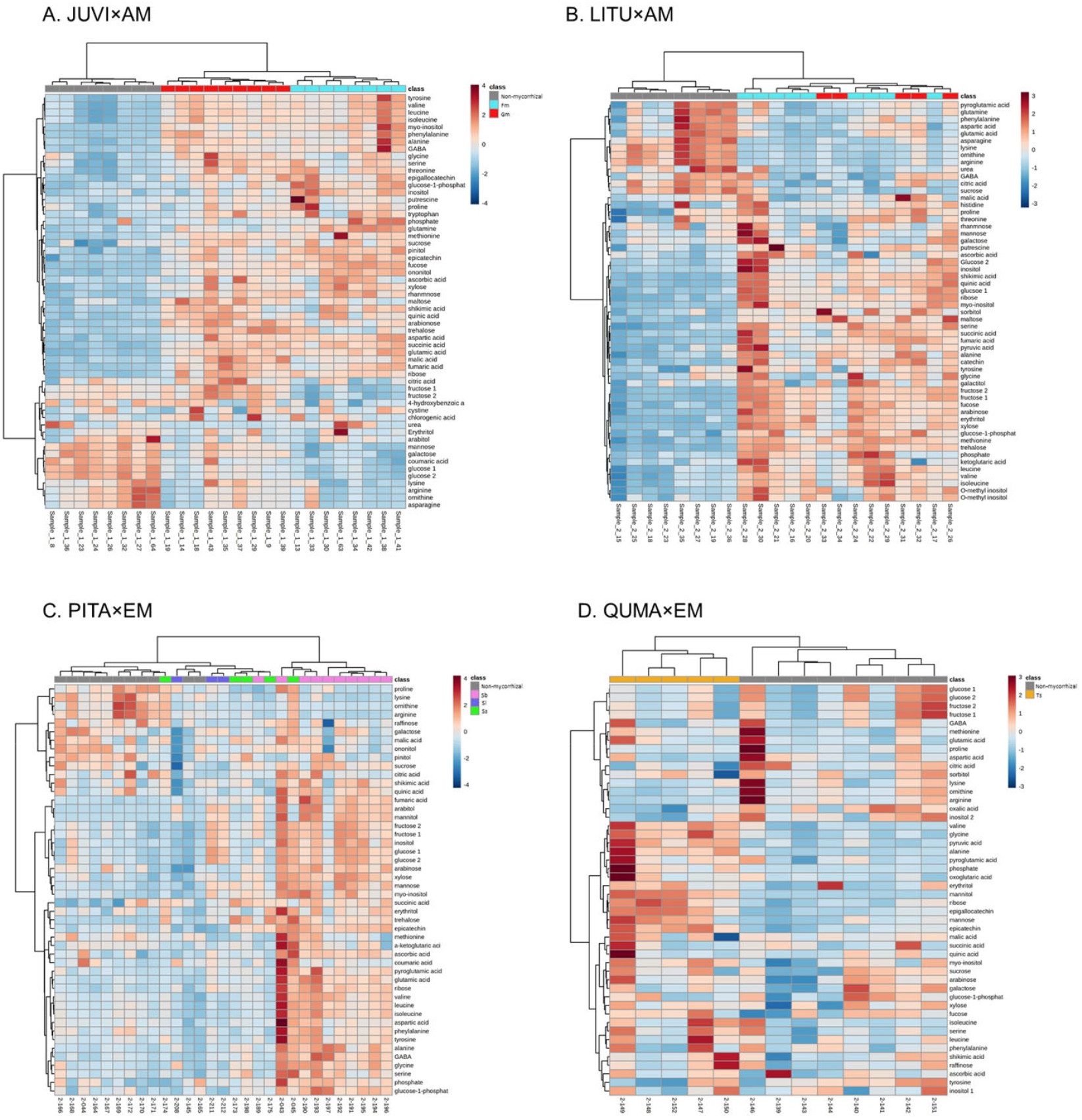
Heatmap and hierarchical clustering analysis of primary metabolites (square root transformed and standardized) in non-mycorrhizal and mycorrhizal roots in eight plant-mycorrhizal fungus combinations across AM and EM lifestyles.

**Fig. S3.**
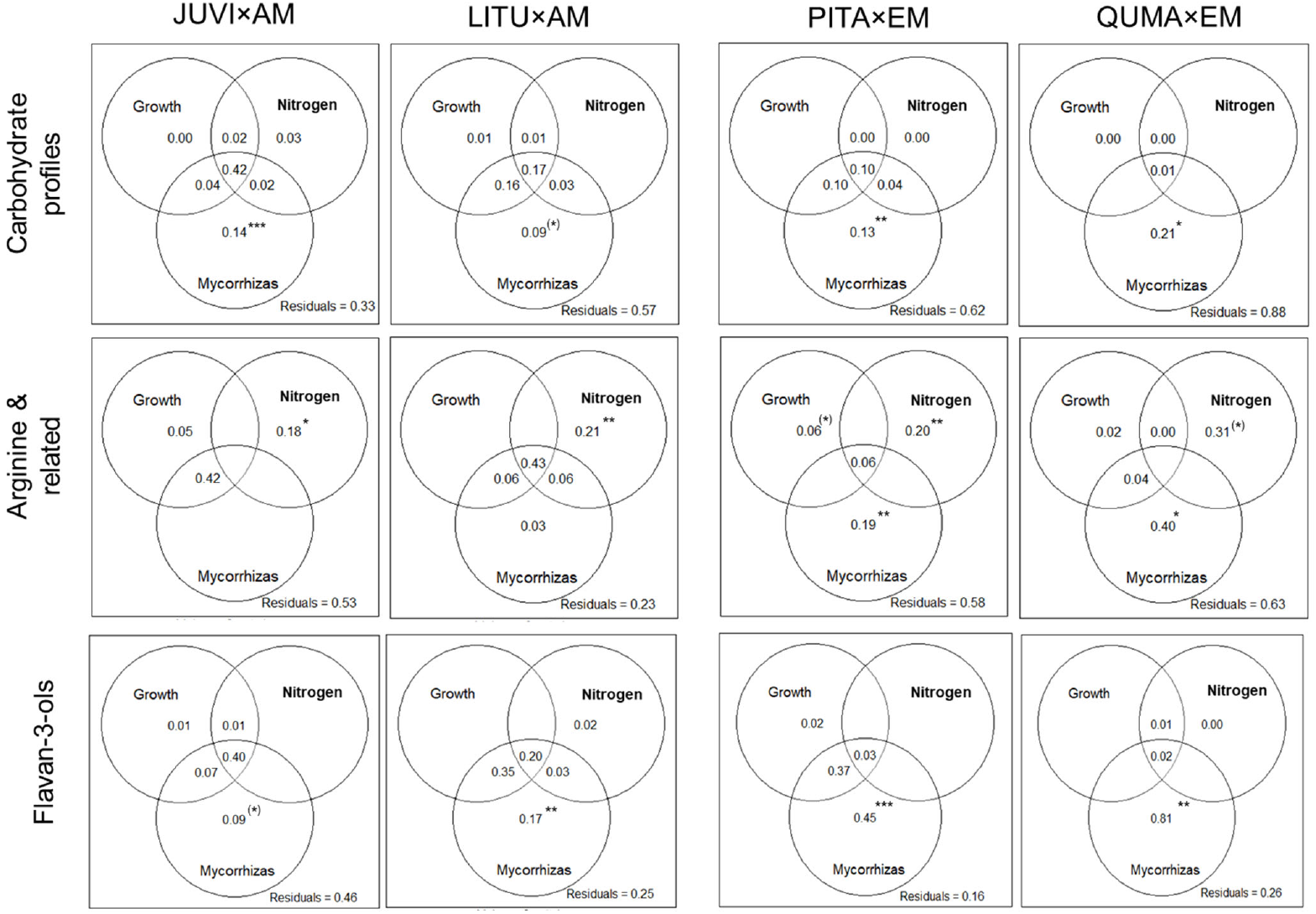
Venn diagram summarizing the variation of carbohydrate profiles, the decrease of arginine & related metabolites, and the accumulation of flavan-3-ols explained by the direct (mycorrhizal colonization) and indirect drivers (growth and tissue N). The parameters that enter the variation partitioning models are listed as follows. Carbohydrate profiles: glucose, fructose, sucrose, polyols, and sugar alcohols (representing >96.8% of total peak area attributed to carbohydrates); arginine & related: arginine and ornithine; flavan-3-ols: summed peak area of all peaks assigned to flavan-3-ols (as shown in Fig. 4B); growth: plant height; nitrogen: nitrogen concentration and the ratio of carbon and nitrogen; mycorrhizas: the presence of each mycorrhizal fungal species. The intersections represent variation that is jointly explained by two or more variable categories and cannot be unambiguously linked to a specific category. Note that the significance of the net effect of each category can be tested statistically but that of the intersections cannot be tested statistically. The significance of net effects of potential drivers was indicated with (*) *P* ≤ 0.1, * *P* ≤ 0.05, ** *P* ≤ 0.01, and *** *P* ≤ 0.001.

**Fig. S4.**
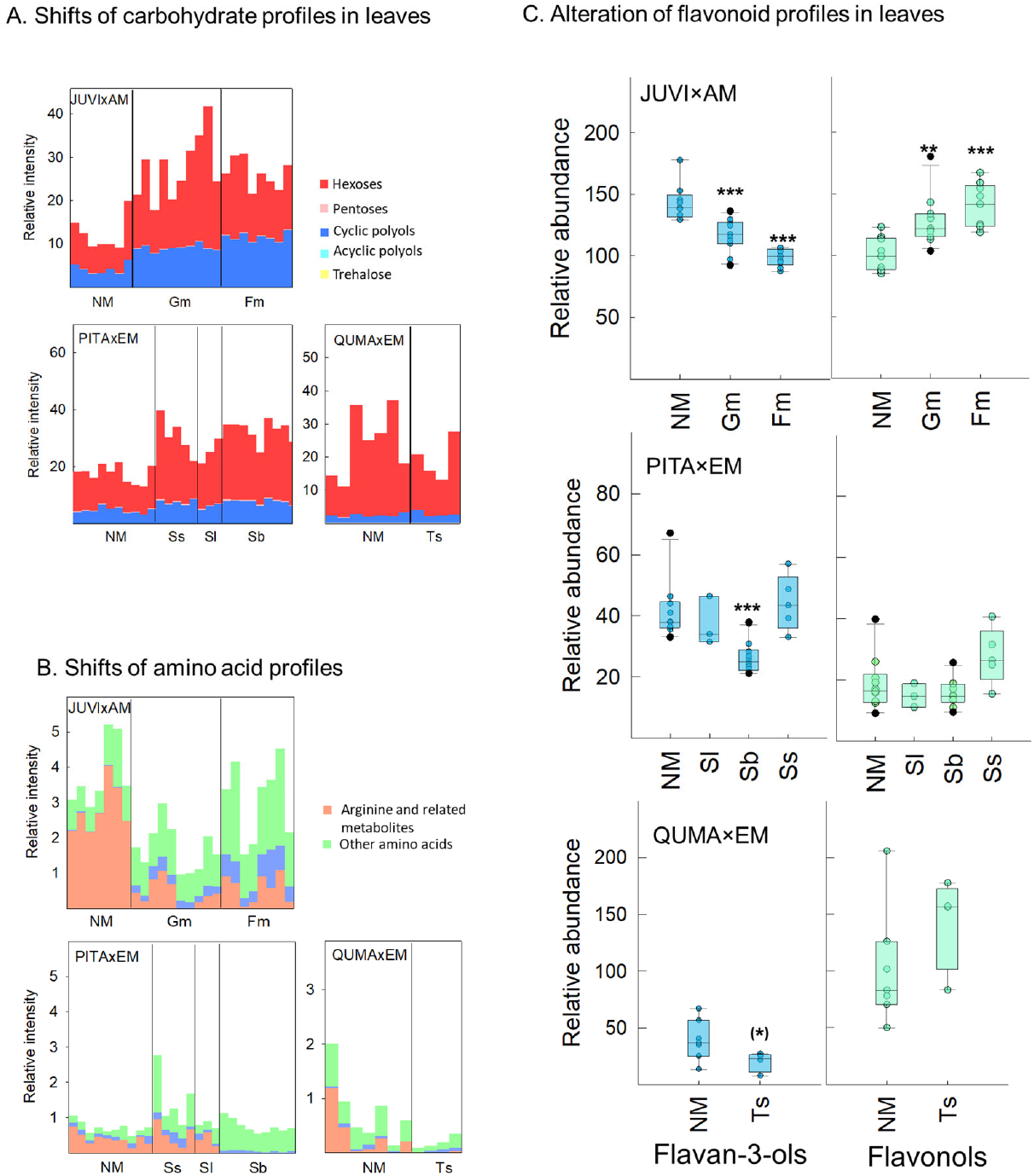
Effects of mycorrhizal associations on leaf metabolites. **A.** The responses of carbohydrate profiles in leaves to mycorrhizal associations did not mirror the accumulation of polyols observed in mycorrhizal-colonized roots. The shifts of carbohydrate profiles were shown by the summed relatively intensity (normalized to the internal standard). **B.** The shifts of amino acid profiles were shown by the summed relatively intensity (normalized to the internal standard) across AM and EM lifestyles. **C.** The responses of flavonoid profiles in leaves to mycorrhizal associations did not mirror the accumulation of flavan-3-ols observed in mycorrhizal-colonized roots. The summed intensity (normalized to internal standards) of all chemical features assigned to flavan-3-ols or flavanols were shown. ^(^*^)^: *P* < 0.1; *: *P* < 0.05; **; *P* < 0.01.***; *P* < 0.001.

**Fig. S5.**
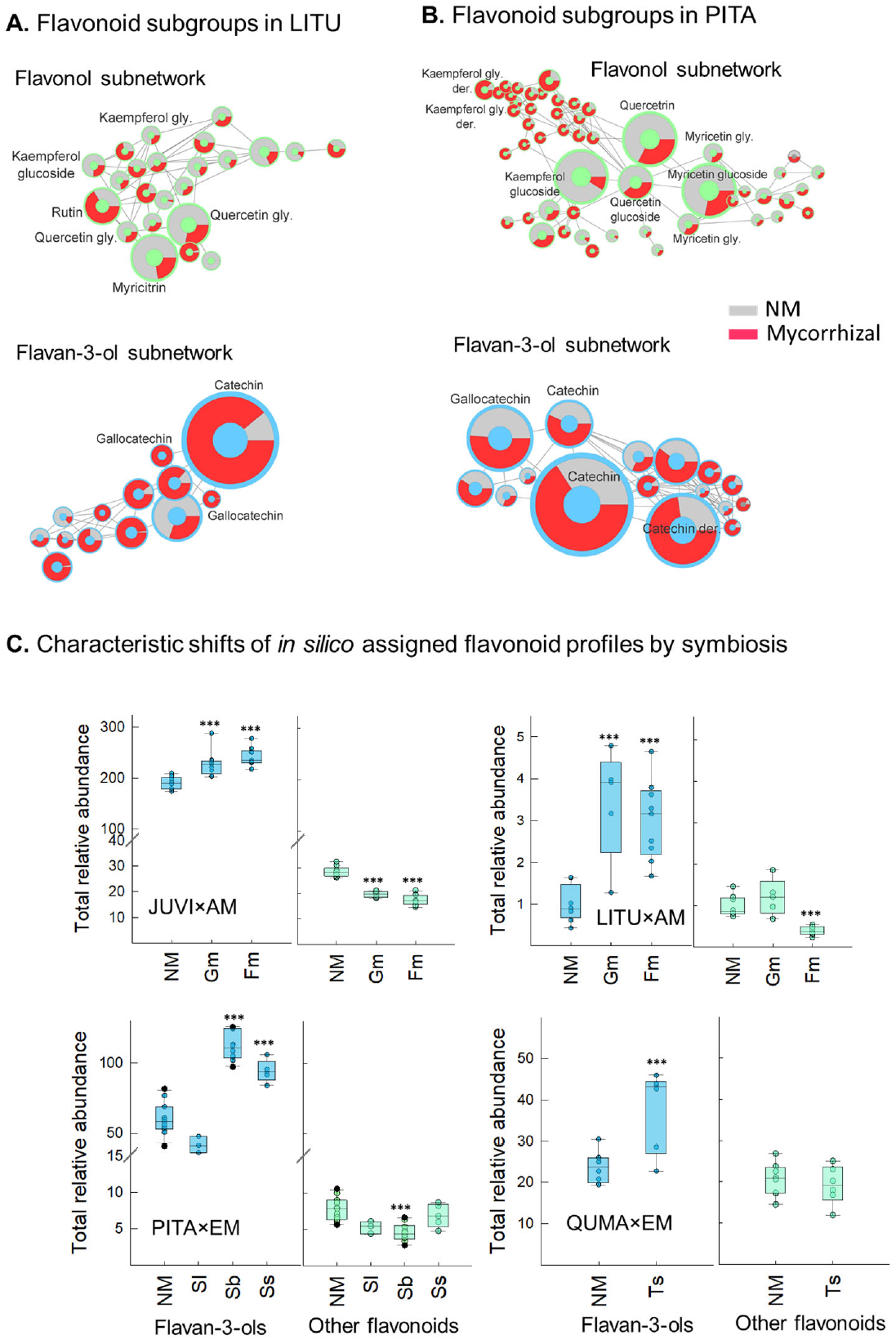
The alterations of *in silico* assigned flavonoid profiles in response to mycorrhizas. The molecular FMNB subnetworks corresponding to two major flavonoid subgroups (*in silico* assigned) that showed differential responses to mycorrhizal colonization were shown for LITU roots colonized by Fm **(A)** and PITA roots colonized by Sb **(B)** as example, with putative annotates shown for several key compounds. The subnetwork in green corresponds to a cluster of glycosylated flavonols (*i.e*., kaempferols, quercetins, and myricetins) and like. The subnetwork in blue corresponds to flavan-3-ols and their derivatives (*i.e*., catechins and gallocatechins). In **(A)** and **(B)**, the comparison between non-mycorrhizal and mycorrhizal roots were shown as rings. The size of rings does not increase linearly with relative intensity but denotes rank relationships among nodes within a subnetwork. der.: derivatives; gly: glycosides. **C.** Characteristic shifts of *in silico* assigned flavonoid profiles in response to mycorrhizal symbiosis featured by the differential changes in the summed intensity (normalized to internal standards) between mass features assigned as flavan-3-ols and those assigned as flavonoids excluding flavan-3-ols. ***: *P* < 0.001.

**Table S1.** The metabolite pathways (KEGG) affected by mycorrhizal symbiosis based on annotated compounds. *P*-values were calculated using Fisher exact tests. (*): *P* < 0.1; *: *P* < 0.05; **: *P* < 0.01; ***: *P* < 0.001.

| Metabolic pathway | JUVI |  | LITU |  | PITA |  |  | QUMA |
| --- | --- | --- | --- | --- | --- | --- | --- | --- |
|  | Fm | Gm | Fm | Gm | Sb | Sl | Ss | Ts |
| Galactose metabolism | * | * | ** | * | (*) |  |  | * |
| Arginine biosynthesis | * | ** | * | ** | * | (*) | * |  |
| Amino sugar and nucleotide sugar metabolism | * | * | ** | ** | * |  |  |  |
| Inositol phosphate metabolism | * | * | * | * |  |  |  |  |
| Starch and sucrose metabolism | * | * |  |  |  |  |  |  |
| Glycolysis/Gluconeogenesis | * | * | (*) | (*) |  |  |  |  |
| Alanine, aspartate and glutamate metabolism | (*) | (*) |  |  |  |  |  | * |
| Pentose phosphate pathway | (*) | (*) |  |  |  | (*) |  |  |
| Arginine and proline metabolism |  | (*) |  |  | (*) |  | (*) |  |
| Citrate cycle |  |  | (*) | (*) |  |  |  |  |
| Fructose and mannose metabolism |  |  |  |  | * | * |  |  |
| 2-Oxocarboxylic acid metabolism |  |  |  |  |  |  | (*) | (*) |
| Flavonoid biosynthesis | * | * | * | * | * |  | (*) | (*) |
| Flavone and flavanol biosynthesis | * | * | * |  | * |  |  |  |
| Biosynthesis of secondary metabolites |  |  | ** | *** |  |  |  |  |
| Phenylpropanoid biosynthesis |  |  | (*) | * |  |  |  |  |

**Table S2** Data file of the details of metabolite annotation (see a separate Excel file).

**Table S3** Data file of annotated compound in each plant species and their respective responses to mycorrhizal colonization. The fold changes of mycorrhizal/non-mycorrhizal for each annotated compound and their *P* values (Student’s *t*-test) are shown.

**Methods S1** Details of the UHPLC-Orbitrap-MS/MS analysis

Samples were separated using an Ultimate 3000 HPLC (Thermo Scientific, Waltham, MA, USA) by an Acquity UPLC HSS T3 column (150 × 2.1 mm, 1.8 μm; Waters Corp., Milford, MA, USA) at 32 °C, and analyzed by an Orbitrap Fusion Tribrid mass spectrometer equipped with an electrospray ion source (Thermo Scientific). The gradient program employed water with 0.1% formic acid as mobile phase A and acetonitrile as mobile phase B, where solvent B increased from 10% to 95% in 12 min and stayed at 95% B for another 5 min, followed by a re-equilibration of 5 min at 10% B. The flow rate was 0.2 ml min^−1^. The mass spectrometer was operated in negative ionization mode with a data-dependent fragmentation (MS2 HCD-CID, Bowers *et al*., 2018) method. The interface settings for the Orbitrap Fusion Tribrid mass spectrometer were: emitter voltage, 3500 V; vaporizer temperature, 325oC; ion transfer tube, 325oC; sheath gas, 55 (Arb); aux gas, 10 (Arb); and sweep gas, 1 (Arb).

## Notes

### Competing Interest Statement

The authors have declared no competing interest.

